# Dropout-Enabled Ensemble Learning for Multi-Scale Biomedical Data

**DOI:** 10.1101/440362

**Authors:** Alexandre Momeni, Marc Thibault, Olivier Gevaert

## Abstract

Leveraging information from multiple scales is crucial to understanding complex diseases such as cancer where this could have a significant impact in improving diagnoses, patient management and treatment decisions. Recent advances in Convolutional Neural Networks (CNNs) have enabled major breakthroughs in biomedical image analysis, in particular for histopathology and radiology images. Our main contribution is a methodology to combine independent CNN models built for these two types of images in order to improve diagnostic accuracy. We train separate CNN models and combine them using a Dropout-Enabled meta-classifier. Our framework achieved second place in the MICCAI 2018 Computational Precision Medicine Challenge.

## 1 Introduction

Recent breakthroughs in deep learning and Convolutional Neural Networks (CNN) have enabled the development of state-of-the art models for many image classification tasks [1,2], with wide applications in areas such as medical image analysis [3,4]. These models have the advantage that they automatically learn the appropriate image features, as opposed to traditional machine learning approaches which require hand-crafted features [5,6]. The features learned by each separate CNN model can also conveniently be stacked together to form the input of a joint model. We follow this approach, building two separate CNNs for radiology and pathology images for the desired task, and subsequently combining them.

Furthermore, the deep learning community has developed many regularization techniques in recent years to deal with the issue of overfitting. This is useful given the scarcity of medical images data. We incorporate these techniques throughout our training process. In particular, we use dropout [7] to generate many variations of the base patients, augmenting the data available for our models. This method lets us leverage the complexity of the medical images, whilst still being able to generalize to unseen data.

Our method achieved second place in the MICCAI 2018 Computational Precision Medicine Challenge, in which participants were asked to classify a cohort of lower grade glioma tumor cases into two sub-types using radiology and histopathology images [8]. Here, we report this model and its results on this classification problem.

## 2 Methodology

### 2.1 Data set

We evaluate our framework on the Computational Precision Medicine Combined Radiology and Pathology classification data set [8]. It contains histopathology and radiology images for 32 glioma patients with annotations for training and validation (16 oligendroglioma and 16 astrocytoma) and 20 glioma patients for testing. These images were collected from several medical centers.

### 2.2 Histopathology image data preprocessing and modeling

Analysis of histopathology slides is a critical step in oncology where it defines the gold standard for diagnosis, prognosis and treatment design. It largely consists of careful microscopic examination of hematoxylin and eosin (H&E) stained tissue sections by a highly skilled pathologist. This can be a tedious, time-consuming and sometimes subjective task. Advances in slide scanning technology and reductions in cost of digital storage capacity have enabled the widespread adoption of digital pathology over the past decade [9]. At the same time, the dramatic increase in computational power and the breakthroughs in deep learning have fueled the rich expansion of visual recognition research [1]. These developments together have led to the rapid emergence of computational histopathology. Most recent works have successfully leveraged state-of-the-art Convolutional Neural Networks (CNNs) for tasks such as disease detection and diagnosis, highlighting the effectiveness and relevance of learned features in complex images such as histopathology slides [10,11,12].

Digital pathology images are massive data sets, which at highest zoom level can have a digital resolution upwards of 100k pixels in both dimensions. However, since localized annotations are very difficult to obtain, data sets may only contain whole slide level diagnosis labels, falling into the category of weakly-supervised learning. To deal with this, we modify existing CNNs architecture to incorporate a Multiple Instance Learning (MIL) framework [13]. This consists in dividing the histopathology slides into small high resolution patches, sampling randomly from these patches and applying patch level CNNs. The MIL framework is then used to combine patch level predictions intelligently and make an overall slide prediction. In the following sub-sections we present our preprocessing pipeline and provide a description of our model for histopathology image analysis.

#### Preprocessing

We perform the following preprocessing steps on the highest slide resolution available:

1. Region of Interest: tissue segmentation is necessary given that there are large areas of white background space in histopathology images which are irrelevant for analysis. We follow a threshold based segmentation method to automatically detect the foreground region. In particular, we first transform the image from RGB to HSV color space and apply Otsu’s method [14] to find the optimal threshold in each channel. The masks are then combined to compute the final tissue segmentation.
2. Tiling: we tile the tissue region extracted from the original slides into 256 × 256 patches.
3. Color Normalization: stain normalization is essential given that the results from the staining procedure can vary greatly. Indeed, differences in slide scanners or staining protocol can materially impact stain color, which in turn can affect algorithm performance. Many methods have been introduced to overcome this well defined problem, including sophisticated end-to-end deep learning solutions [15]. For simplicity, we resort to a histogram equalization algorithm as proposed in [16].

#### CNN model

Convolutional Neural Networks (CNNs) are very computationally expensive to train in practice and they require a large data set to avoid overfitting. We used the DenseNet-169 [17] architecture starting with initial pretrained parameters from ImageNet [18] and we used fine tuning of these parameters to speed up convergence. We also replaced the last fully connected layers to be compatible with our classification task, using dropout for regularization. Our CNN is trained at the patch level and then produces an average slide level score by averaging all sampled patches for a given patient. The steps of the model are described schematically in Figure 1.

**Fig. 1.**
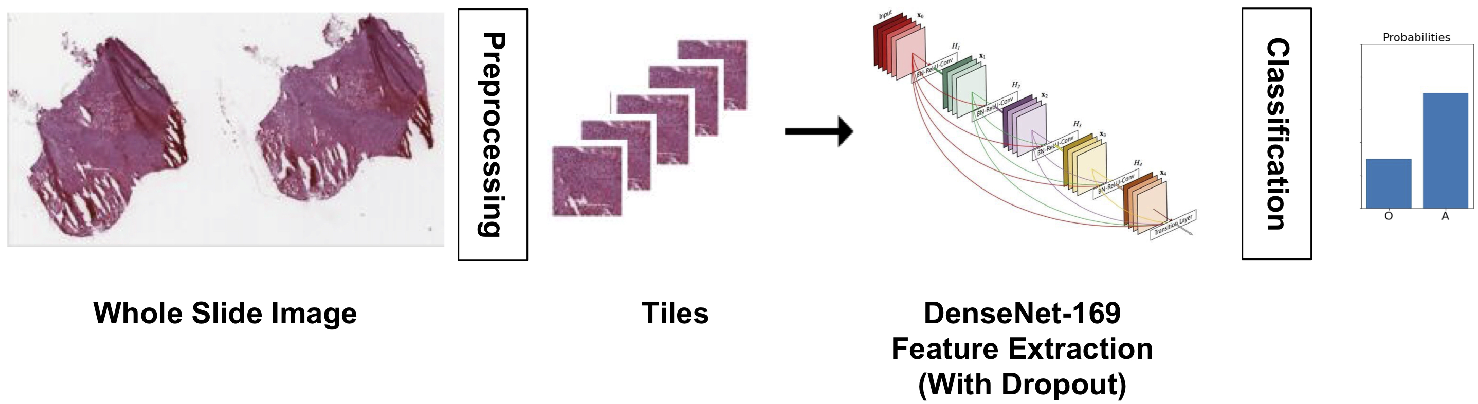
Description of the histopathology prediction pipeline. The slides are tiles, features are extracted with a finetuned DenseNet, classification is made by a final fully connected network.

### 2.3 Radiology image data preprocessing and modeling

Our computational task consists in evaluating also the radiology images of a patient’s lower grade glioma in the form of multi-modal Magnetic Resonance (MR) images. An automated algorithm for classification should analyze these 3D MR images, aggregate local information to understand where the tumor is located, and then compute metrics on these tumors (e.g. capturing size, intensity or texture). These features would then have to be evaluated so that a final decision rule can be devised.

A classical approach to this problem would be to compute hand-made features from the segmentation mask of the tumor [19,20] - this segmentation mask being obtainable in a completely computational fashion - and then to use these features to classify the patient’s status using a linear classification algorithm in the feature space [4,20]. However, this method would have to rely on complex feature extraction methods [21], mimicking a radiologist’s analysis, which can be time-consuming and difficult to hard-code.

Convolutional Neural Networks (CNNs), on the other hand, have proven effective in reaching state-of-the art results in computational analysis of biomedical images for disease detection and diagnosis ([22,2]). Consequently, a newer, more data-driven way to tackle this classification task would be to rely on CNNs for feature extraction and image downsampling, letting us train a neural network through backpropagation to perform the classification task. This CNN can take into input either the raw image channels (i.e. MR image modalities), or the tumor segmentation mask, or both. However, in this contribution, we use the raw modalities as inputs, in order not to rely on external data, and to evaluate the stand-alone performance of our approach.

#### Preprocessing

We applied preprocessing steps which have been used for the tumor segmentation task, from brain MR images. It consists in the tasks of bias correction, skull-stripping, and registration:

1. Bias Correction: we used the **`FSL`** library to remove the bias fields from all modality images, as provided by [23];
2. Skull Removal: we used the Brain Extraction Tool (BET, [24]) from the same library to remove the skull structure from these images;
3. Co-registration: we used the **`reg_aladin`** command from the **`niftyreg`** library to co-register the modalities on the same standard grid [25].

We also enriched our data by randomly flipping scans along the x-axis at train time.

Furthermore, we chose to only rely on the contrast-enhanced T1 and FLAIR MR modalities for this radiology task. Indeed, T1+contrast is sufficient to magnify the grey matter; while the FLAIR modality lets the unusual brain structures appear explicitly. This resulted in 27 patients out of 32 available in the train set. The images were then re-sized to 320×320×24.

#### CNN model

The input of this model is a 3-D voxel image with three spatial dimensions times two modalities per voxel, the FLAIR and T1+contrast MR images. We used an 8-layers 3-D CNN to extract deep features from the images, with three 3-D maxpooling operations to reduce the sample size. The convolution layers have a receptive field of five cells from the previous layers, and extract from eight features on the first layer, to 64 channels on the last layer. The maxpooling layers downsample the images by a factor of two in each dimension.

After the last convolutional layer is applied, we averaged the extracted features over all the 3D space to have a unique 100-dimensional feature vector for the patient. This feature vector is then connected to a 1-dimensional output for a classification task with cross-entropy loss, and the whole network is trained with the back-propagation algorithm.

We inserted a total of 8 dropout layers throughout the network, both to avoid overfitting, and, most importantly for the ensemble learning step, to let us evaluate the network at test time. The overall pipeline is sketched in Figure 2.

**Fig. 2.**
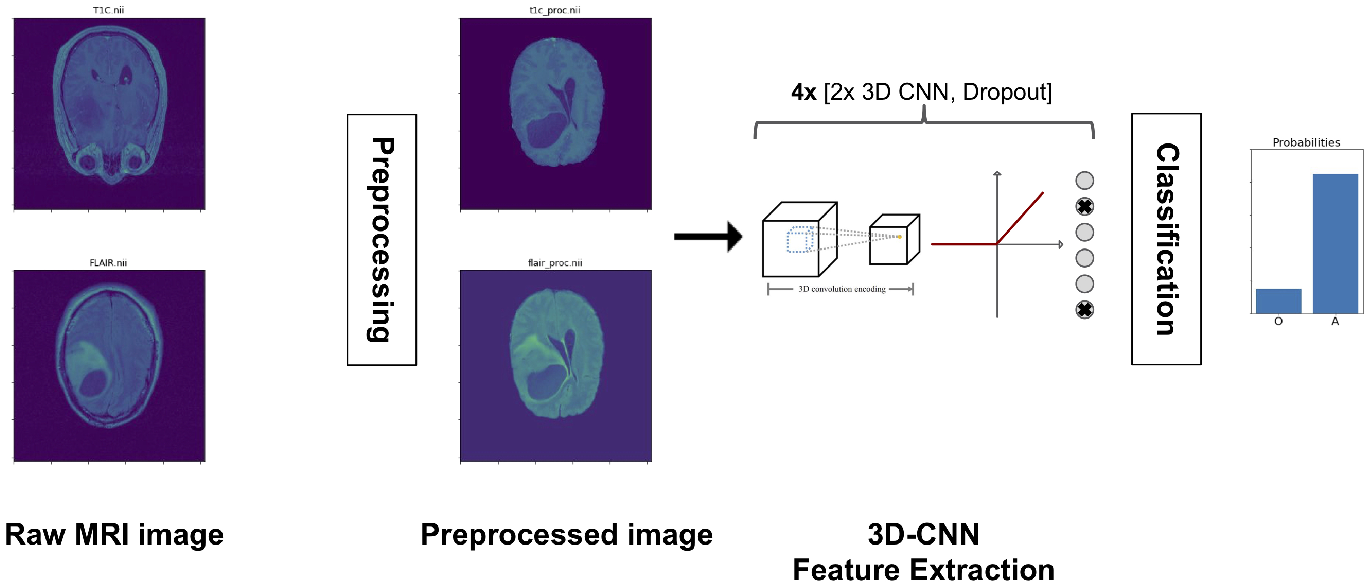
Description of the radiology prediction pipeline. Preprocessing removes the skull and balances the channels, features are extracted with a 3D-CNN, classification is made with a fully connected network.

### 2.4 Meta-classifier methodology image data preprocessing and modeling

Next, to combine the two CNNs modeling the histopathology and MR images, we introduce a meta-algorithm to combine the individual models. Our ensemble learning methodology allows each model to be trained separately, and combines their predictions into a single, more robust output.

#### Meta-classifier for models combination

Since these models rely on very different data sources, with different scales, batching methods and actual biological meaning, it is not always possible to train an end-to-end backpropagation algorithm combining these. A crucial limitation is the computational power required for these models to run simultaneously, especially in terms of storage space. Another inconvenience to the co-training the models is reduced modularity, where a big advantage is if more data from an extra level of biomedical data can be added without having to reconstruct and train the other models.

Our individually trained networks provide us with classification scores for each patient in our data set, quantifying the learned probability that the patient belongs to one category or the other. We then concatenate these individual scores into a two-dimensional vector and train a meta-classifier on these vectors of scores to make a combined prediction for the patient. The meta-classifier using information from both models is a classical classification algorithm, in this case we used a random forest. In order to quantify our ability to fit a meta-classifier, as well as its ability to generalize to unseen data, we used four-fold cross-validation combined with a random forest classifier with ten trees.

#### Dropout-enabled scores consolidate the models’ outputs

Next, we evaluated the use of regularization through dropout for the ensemble learning phase. The idea is to activate the models’ random dropout layers at score extraction time, so that individual models produce multiple classification scores for each patient. This array of scores is then averaged into a global patient score. This approach is comparable to the reasoning of [26], where it is applied to RNNs for final classification, while we use it as the penultimate step for our classifier combination.

The rationale behind this choice is that individual classification CNNs were trained with dropout, such that they have learned several robust ways to classify a patient’s status. Consequently, scores extracted with dropout capture the variability of the models’ prediction. Averaging these scores into a global dropout-enabled patient score removes the noise which emerges when running a single score prediction. It yields more robust aggregated classification scores which leverages the networks’ structure, without the need of an end-to-end joint training.

We quantified the extent to which this technique improves the class separation, by running the same analysis as before. We evaluated the performance of a random forest classifier using four-fold cross-validation on the dropout-enabled scores. We used accuracy and Area Under the ROC curve (AUC) to evaluate the models.

## 3 Results

Figure 3.a) shows the output classification scores of each patient, respectively from the pathology and the radiology models, depending on their status, on the training set. This shows that these scores let us define with some confidence some of the patients’ status. However, there does not seem to be a simple separation between the two classes leading to a reliable meta-classifier.

Next, Table 1 presents the classification results of this method on the raw scores, as estimated via vross-validation. It appears that the pathology model is a good predictor of the patients’ statuses (accuracy 78%, AUC 0.83); while radiology does not make a good contribution (accuracy 53%, AUC 0.54). The combined model performed well, but not significantly better than the pathology model, suggesting that the radiology model does not contribute to the performance (accuracy 81%, AUC 0.84).

**Fig. 3.**
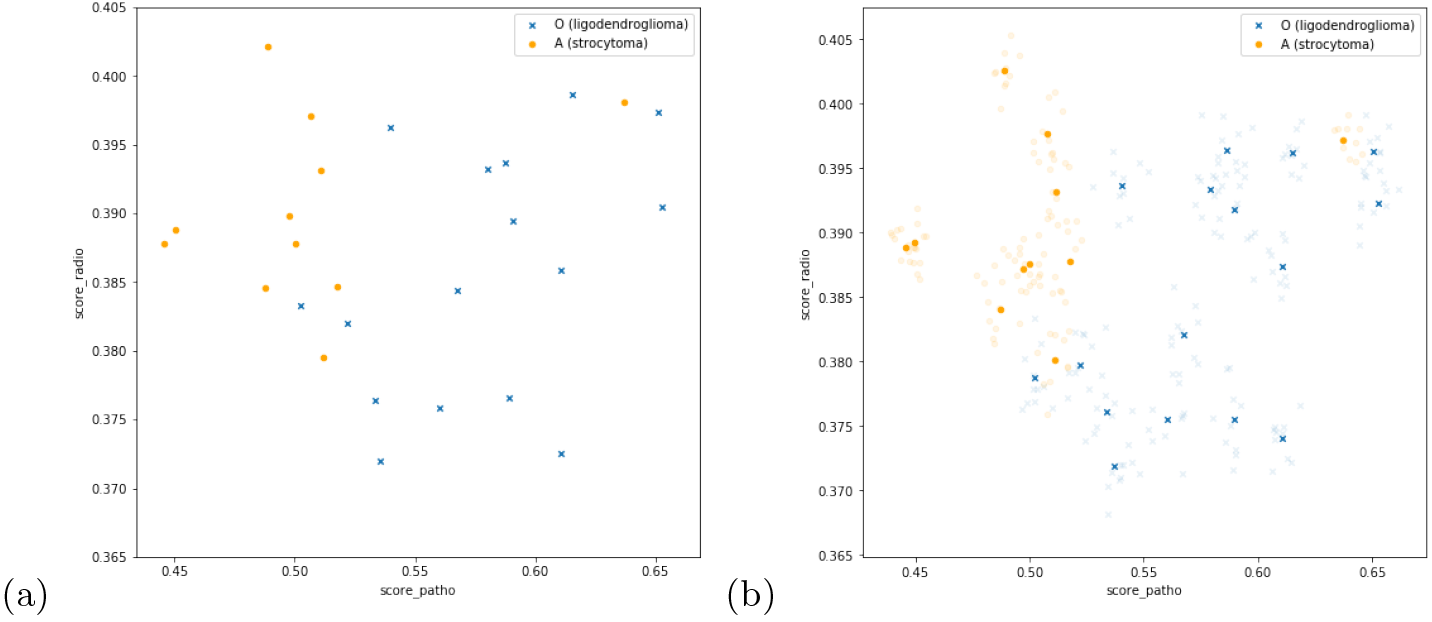
Scatter plot of the patients classification scores, according to the pathology model on the x-axis, and the radiology model on the y-axis, colored by their status. **(a)**: Raw classification scores. **(b)**: Dropout-Enabled classification scores; the patient-averaged scores are highlighted.

**Table 1.** Accuracy and AUC for classifying subtype of brain tumors using 4-fold crossvalidation and a random forest classifier, both for raw scores and average dropout-enabled scores.

| Model | Data | ACCURACY | AUC |
| --- | --- | --- | --- |
| Raw scores | Radiology | 53% | 0.54 |
|  | Histopathology | 78% | 0.83 |
|  | Combined | 81% | 0.84 |
| Dropout-Enabled scores | Radiology | 70% | 0.70 |
|  | Histopathology | 79% | 0.83 |
|  | **Combined** | **85%** | **0.92** |

Next, Figure 3.b) presents the classification scores of the patients of both classes, after the dropout-enabled aggregation phase has been applied. This shows that the patients which were in grey zones are now easier to classify, resulting in more separable classes. The performances of the dropout-enabled scores are summarized in Table 1. The classification scores of the histopathology model alone are similar to the previous ones (accuracy 79%, AUC 0.83). Most importantly, we see that this technique improves the performance of the radiology-based model (accuracy 70%, AUC 0.70), and contributes to enhancing the combined classifier (accuracy 85%, AUC 0.92).

## 4 Conclusion

We have proposed a new approach to combine individual CNN classifiers leveraging dropout to combine two types of biomedical image data: histopathology and radiology images, for classification. We avoided training an end-to-end CNN training through backpropagation on heterogeneous data, as this would reduce modularity and be computationally expensive. Our method allows us to make more relevant decisions for outlier cases, based on separate predictions. Next, using dropout to generate several predictions of a patient’s data can be used in further directions, that will be studied independently in further works: we plan on using the same analysis at feature-level instead of score-level, which would allow us to analyze the non-linear interactions between multi-scale data. Finally, applying this technique to glioma subtype classification using radiology and histopathology images provides initial evidence that a multi-scale model can improve the prediction of a patient’s diagnosis. Extending this analysis to bigger cohorts and other classification problems is warranted and will allow to study the multi-scale model in more detail.

## 5 Acknowledgements

Research reported in this publication was supported by the National Institute of Biomedical Imaging and Bioengineering of the National Institutes of Health under Award Number R01EB020527. The content is solely the responsibility of the authors and does not necessarily represent the official views of the National Institutes of Health.

## References

1. Yann LeCun, Yoshua Bengio, and Geoffrey Hinton. Deep learning. nature, 521(7553):436, 2015.

2. Andre Esteva, Brett Kuprel, Roberto A Novoa, Justin Ko, Susan M Swetter, Helen M Blau, and Sebastian Thrun. Dermatologist-level classification of skin cancer with deep neural networks. Nature, 542(7639):115, 2017.

3. Justin Ker, Lipo Wang, Jai Rao, and Tchoyoson Lim. Deep learning applications in medical image analysis. IEEE Access, 6:9375–9389, 2018.

4. Darvin Yi, Mu Zhou, Zhao Chen, and Olivier Gevaert. 3-d convolutional neural networks for glioblastoma segmentation. arXiv preprint arXiv:1611.04534, 2016.

5. Olivier Gevaert, Sebastian Echegaray, Amanda Khuong, Chuong D Hoang, Joseph B Shrager, Kirstin C Jensen, Gerald J Berry, H Henry Guo, Charles Lau, Sylvia K Plevritis, et al. Predictive radiogenomics modeling of egfr mutation status in lung cancer. Scientific reports, 7:41674, 2017.

6. Shaimaa H Bakr, Sebastian Echegaray, Rajesh P Shah, Aya Kamaya, John Louie, Sandy Napel, Nishita Kothary, and Olivier Gevaert. Noninvasive radiomics signature based on quantitative analysis of computed tomography images as a surrogate for microvascular invasion in hepatocellular carcinoma: a pilot study. Journal of Medical Imaging, 4(4):041303, 2017.

7. Nitish Srivastava, Geoffrey Hinton, Alex Krizhevsky, Ilya Sutskever, and Ruslan Salakhutdinov. Dropout: a simple way to prevent neural networks from overfitting. The Journal of Machine Learning Research, 15(1):1929–1958, 2014.

8. Miccai cpm competition.

9. Anant Madabhushi and George Lee. Image analysis and machine learning in digital pathology: Challenges and opportunities, 2016.

10. Dan C Cireşan, Alessandro Giusti, Luca M Gambardella, and Jürgen Schmidhuber. Mitosis detection in breast cancer histology images with deep neural networks. In International Conference on Medical Image Computing and Computer-assisted Intervention, pages 411–418. Springer, 2013.

11. Yun Liu, Krishna Gadepalli, Mohammad Norouzi, George E Dahl, Timo Kohlberger, Aleksey Boyko, Subhashini Venugopalan, Aleksei Timofeev, Philip Q Nelson, Greg S Corrado, et al. Detecting cancer metastases on gigapixel pathology images. arXiv preprint arXiv:1703.02442, 2017.

12. Le Hou, Dimitris Samaras, Tahsin M Kurc, Yi Gao, James E Davis, and Joel H Saltz. Patch-based convolutional neural network for whole slide tissue image classification. In Proceedings of the IEEE Conference on Computer Vision and Pattern Recognition, pages 2424–2433, 2016.

13. Thomas G Dietterich, Richard H Lathrop, and Tomás Lozano-Pérez. Solving the multiple instance problem with axis-parallel rectangles. Artificial intelligence, 89(1-2):31–71, 1997.

14. Nobuyuki Otsu. A threshold selection method from gray-level histograms. IEEE transactions on systems, man, and cybernetics, 9(1):62–66, 1979.

15. M Tarek Shaban, Christoph Baur, Nassir Navab, and Shadi Albarqouni. Staingan: Stain style transfer for digital histological images. arXiv preprint arXiv:1804.01601, 2018.

16. Denis Nikitenko, Michael Wirth, and Kataline Trudel. Applicability of white-balancing algorithms to restoring faded colour slides: An empirical evaluation. Journal of Multimedia, 3(5), 2008.

17. Gao Huang, Zhuang Liu, Laurens Van Der Maaten, and Kilian Q Weinberger. Densely connected convolutional networks. In CVPR, volume 1, page 3, 2017.

18. Jia Deng, Wei Dong, Richard Socher, Li-Jia Li, Kai Li, and Li Fei-Fei. Imagenet: A large-scale hierarchical image database. In Computer Vision and Pattern Recognition, 2009. CVPR 2009. IEEE Conference on, pages 248–255. Ieee, 2009.

19. Shuo Wang, Mu Zhou, Zaiyi Liu, Zhenyu Liu, Dongsheng Gu, Yali Zang, Di Dong, Olivier Gevaert, and Jie Tian. Central focused convolutional neural networks: Developing a data-driven model for lung nodule segmentation. Medical image analysis, 40:172–183, 2017.

20. Sebastian Echegaray, Olivier Gevaert, Rajesh Shah, Aya Kamaya, John Louie, Nishita Kothary, and Sandy Napel. Core samples for radiomics features that are insensitive to tumor segmentation: method and pilot study using ct images of hepatocellular carcinoma. Journal of Medical Imaging, 2(4):041011, 2015.

21. Haruka Itakura, Achal S Achrol, Lex A Mitchell, Joshua J Loya, Tiffany Liu, Erick M Westbroek, Abdullah H Feroze, Scott Rodriguez, Sebastian Echegaray, Tej D Azad, et al. Magnetic resonance image features identify glioblastoma phenotypic subtypes with distinct molecular pathway activities. Science translational medicine, 7(303):303ra138–303ra138, 2015.

22. Zeynettin Akkus, Alfiia Galimzianova, Assaf Hoogi, Daniel L Rubin, and Bradley J Erickson. Deep learning for brain mri segmentation: state of the art and future directions. Journal of digital imaging, 30(4):449–459, 2017.

23. Yongyue Zhang, Michael Brady, and Stephen Smith. Segmentation of brain mr images through a hidden markov random field model and the expectation-maximization algorithm. IEEE transactions on medical imaging, 20(1):45–57, 2001.

24. Stephen M Smith. Fast robust automated brain extraction. Human brain mapping, 17(3):143–155, 2002.

25. Sébastien Ourselin, Alexis Roche, Gérard Subsol, Xavier Pennec, and Nicholas Ayache. Reconstructing a 3d structure from serial histological sections. Image and vision computing, 19(1-2):25–31, 2001.

26. Yarin Gal and Zoubin Ghahramani. A theoretically grounded application of dropout in recurrent neural networks. In Advances in neural information processing systems, pages 1019–1027, 2016.

